## Supplementary material for "High-molecular-weight polymers from dietary fiber drive aggregation of particulates in the murine small intestine"

**Figure 1 – figure supplement 1, Figure 4 – figure supplements 1-3, Figure 5 – figure supplement 1, Figure 6 – figure supplements 1-3**

**Tables 1 – 8**

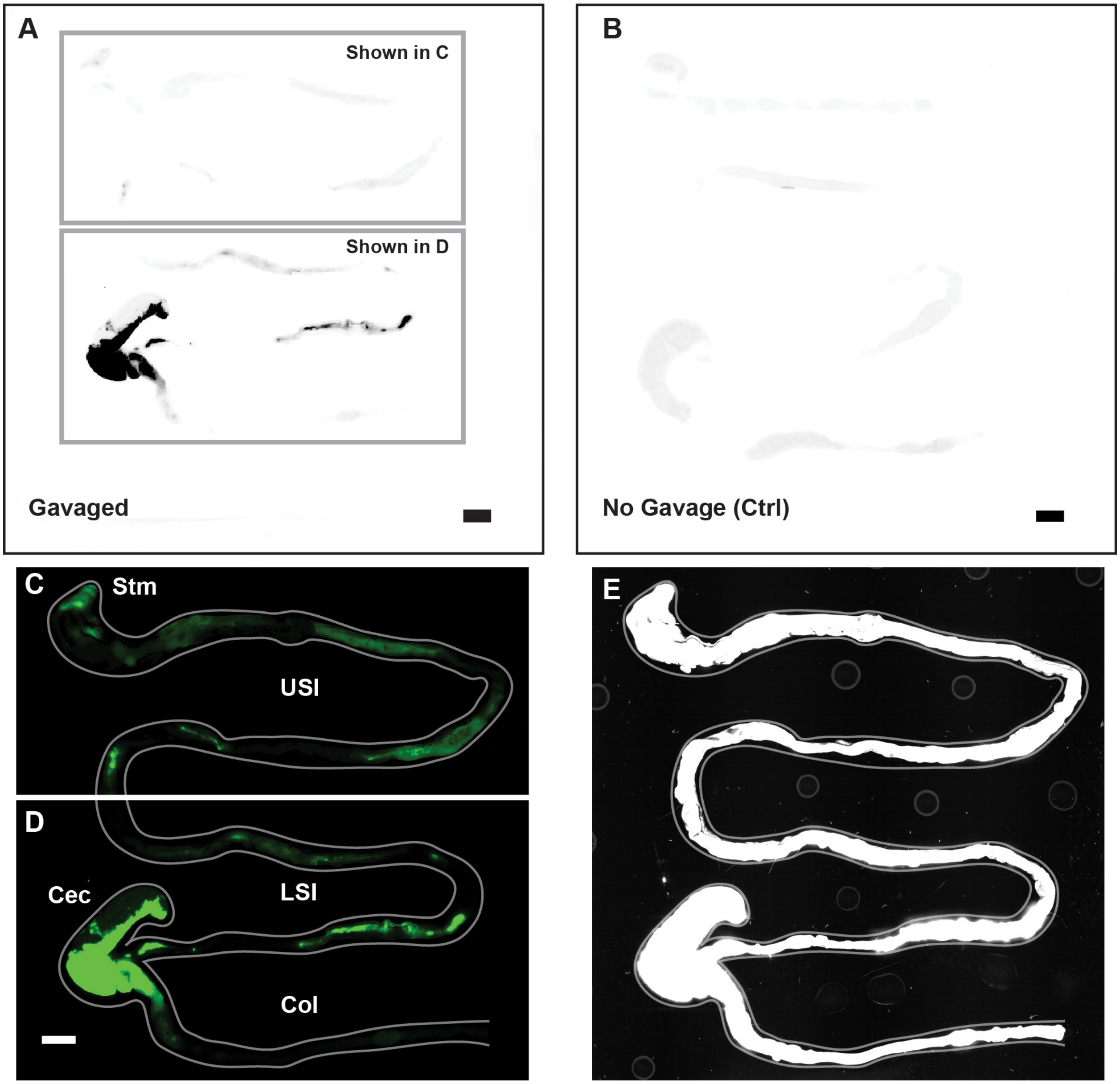


**Figure 1 – figure supplement 1.** Overview of image processing for fluorescent scanner images appearing in Figure 1. (A) Unmodified fluorescent scanner images of the gastrointestinal tract of a mouse gavaged with 1 µm-diameter PEG-coated particles (prior to the contrast and color-adjustments shown in Fig. 1A–B). Scale bar is 0.5 cm. Boxes indicate the regions that are shown in panels C and D. (B) Unmodified fluorescent scanner image of the gut of a mouse that has not been gavaged with particles. Scale bar is 0.5 cm. (C and D). The contrast and color-adjusted images that appear in Fig. 1A–B. (E) Contrast-adjusted image of Figures 1A-B that was used to trace the outline of the gut shown in Fig. 1A–B (and panel C and D of this figure). Outline of gut is shown in grey on both C, D, and E.

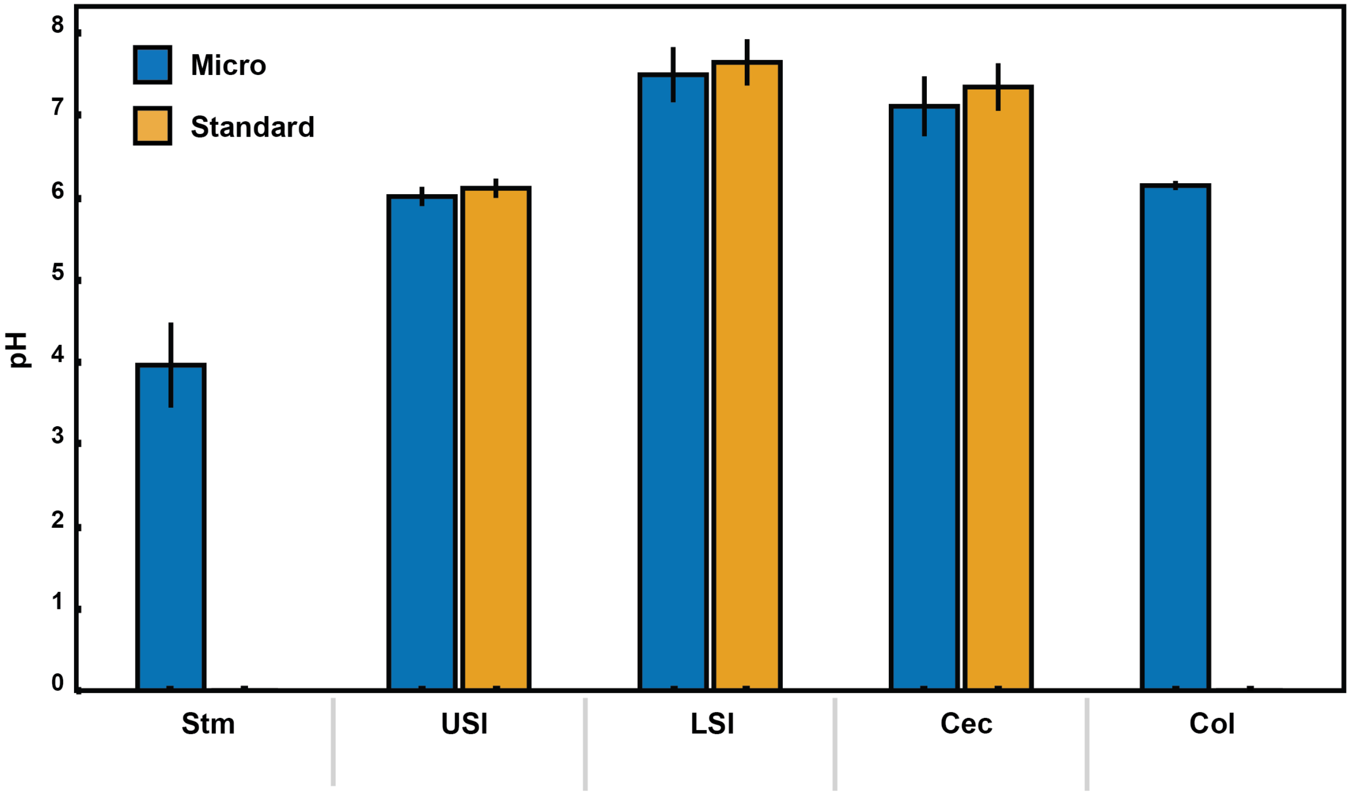


**Figure 4 – figure supplement 1**. pH measurements of luminal fluid from different sections of the gastrointestinal tract. Measurements were conducted on pooled samples of luminal fluid collected from two groups of mice. Each measurement was repeated three times, and the error bars are the standard deviation across the six trials (three trials per group). Micro (blue) indicates measurements that were conducted using a micro pH electrode. Standard (orange) indicates measurements that were conducted using a standard pH electrode. For the stomach and colon samples there was insufficient luminal fluid from both groups to submerge the tip of the standard pH electrode, so measurements were only taken with the micro pH electrode. Stm = stomach, USI = upper small intestine. LSI = lower small intestine, Cec = cecum, and Col = colon.

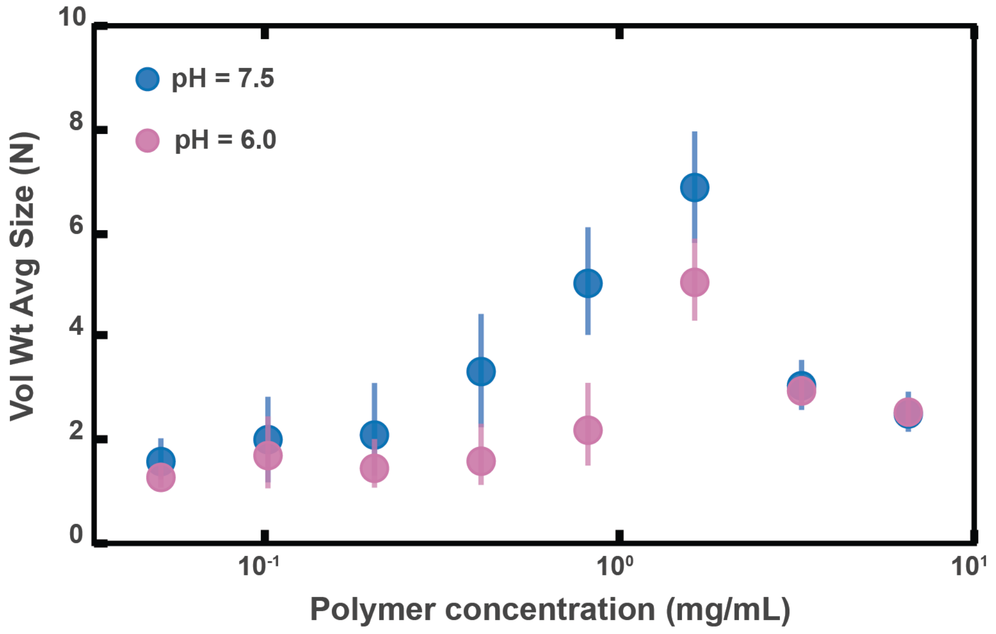


**Fig. 4 – figure supplement 2. Aggregation of PEG-coated particles in model polymer solutions with different pH (A)** Volume-weighted average sizes for serial dilutions of 1 MDa PEG solutions in a phosphate buffered saline solution with $pH = 6.0 \pm 0.1$ (labeled pH = 6.0) and in Hank's balanced salt solution (HBSS) with $pH = 7.6 \pm 0.1$ (same data from Figure 4D). Volume-weighted average sizes are plotted on the vertical axis in terms of number of particles per aggregate (N) against polymer mass concentration ($c_p$) in mg/mL. The vertical error bars are 95% empirical bootstrap CI (see *Materials and Methods* for bootstrapping procedure).

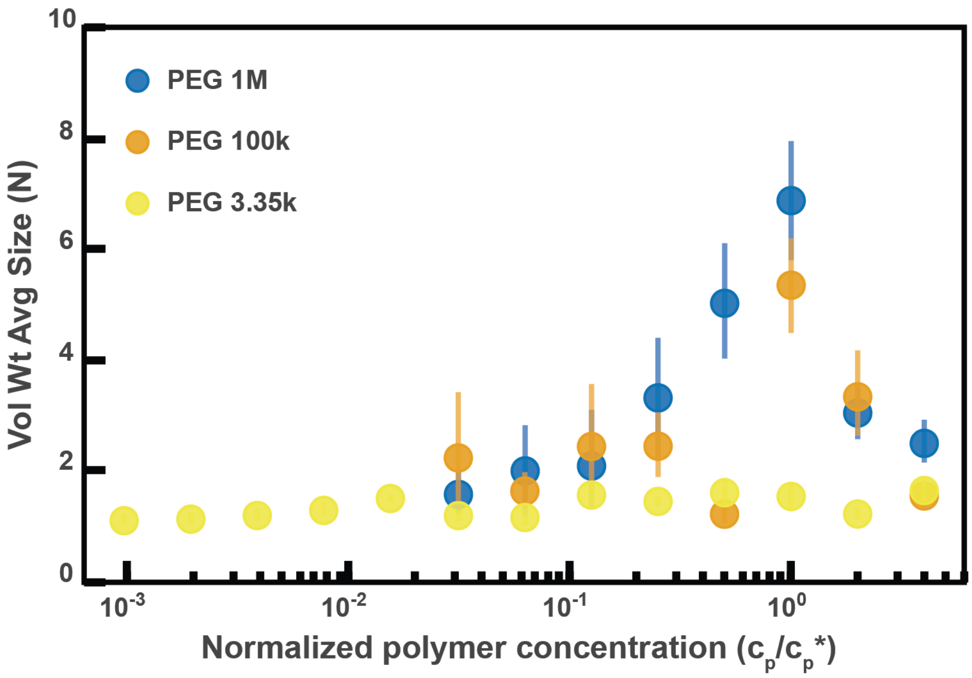


**Fig. 4 – figure supplement 3. Aggregation of PEG-coated particles in model polymer solutions from Figure 4D normalized by polymer overlap concentration.** Volume-weighted average sizes for serial dilutions of 1 MDa PEG solutions in Hank's balanced salt solution (HBSS). Volume-weighted average sizes are plotted on the vertical axis in units of number of particles per aggregate (N) against the "normalized polymer concentration." The normalized polymer concentration is the polymer mass concentration ($c_p$) in mg/mL divided by the overlap concentration of each polymer solution ($c_p^*$) in mg/mL. The overlap concentrations for PEG 1 MDa, 100 kDa, and 3350 Da are $c_p^*$ = 1.6, 8.6, and 52.6 mg/mL, respectively. The vertical error bars are 95% empirical bootstrap CI (see *Materials and Methods* for bootstrapping procedure).

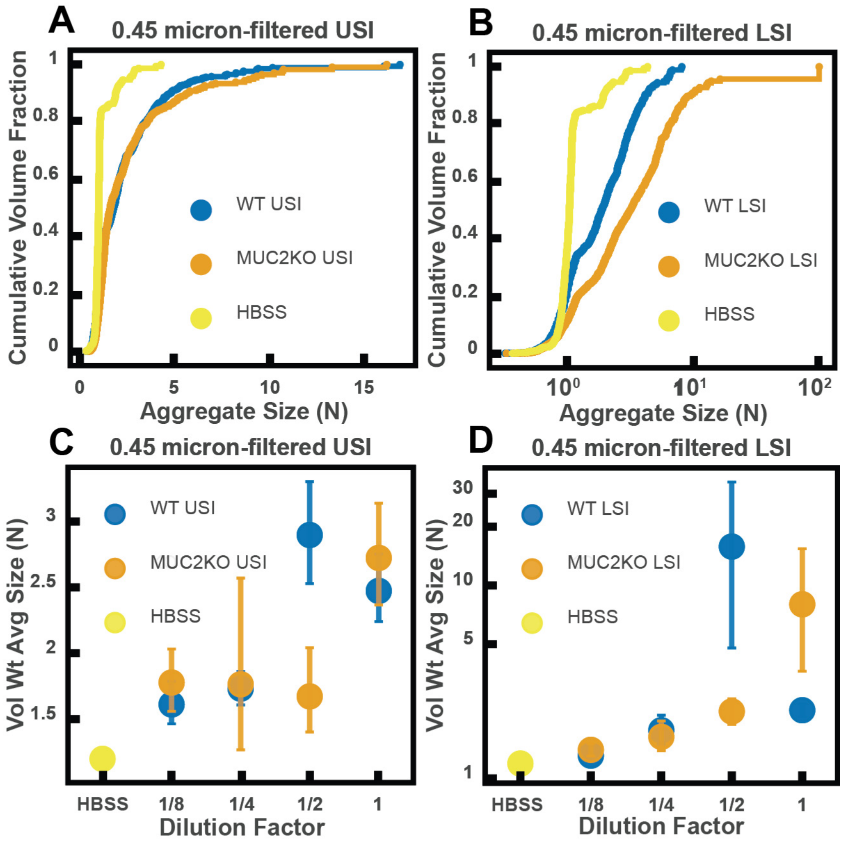


**Figure 5 – figure supplement 1.** *Ex vivo* aggregation in 0.45 μm-filtered luminal fluid from the small intestines (SI) of wild-type (WT) and MUC2 knockout (MUC2KO) mice. (**A** and **B**) Volume-weighted empirical cumulative distribution functions (ECDFs) comparing aggregation of the particles in undiluted, 0.45-μm-filtered samples from the upper (A) and lower (B ) SI of two separate groups of WT and MUC2KO mice to the control (particles suspended in HBSS). The vertical axis is the cumulative volume fraction of the total number of particles in solution in an aggregate of a given size. The horizontal axis is aggregate size in number of particles per aggregate (N). (C and D) Volume-weighted average aggregate sizes (Vol Wt Avg Size) for serial dilutions of 0.45 μm-filtered samples from the upper (C) and lower (D) SI of two separate groups of WT and MUC2KO mice. Volume-weighted average sizes are plotted on the vertical axis in terms of number of particles per aggregate (N). The dilution factor is plotted on the horizontal axis, where a dilution factor of 1 is undiluted and ½ is a two-fold dilution. The control (particles suspended in HBSS) is plotted as a dilution factor of 0. The vertical error bars are 95% empirical bootstrap CI using the bootstrapping procedure described in *Materials and Methods*.

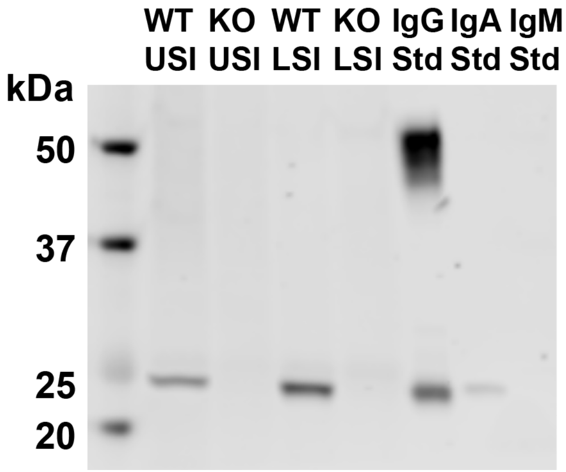


**Figure 6 – figure supplement 1.** Western blots of 30 µm-filtered samples from the small intestine (SI) of wild-type (WT) and Rag1 knockout (Rag1KO) mice. WT USI = WT upper SI; KO USI = KO lower SI; WT LSI = WT lower SI; KO USI = KO upper SI. For the detection of IgG, 1:10,000 dilutions of Li-Cor IRDye 800 CW Goat Anti-Mouse IgG was used. Because the Anti-IgG antibody appears to be binding to just the light chains (around 25 kDa), we suspect that it is mostly binding to IgA. Li-Cor's published validation (https://www.licor.com/bio/products/reagents/secondary_antibodies/irdye_800cw.html) found that the antibody binds to the heavy and light chains of IgG and just the light chains of IgA. Because we see binding of the antibody to both the heavy and light chains in the IgG standard, but only binding to a light chain in the SI samples and the IgA control, this suggests that we are detecting the light chains of IgA in the SI samples.

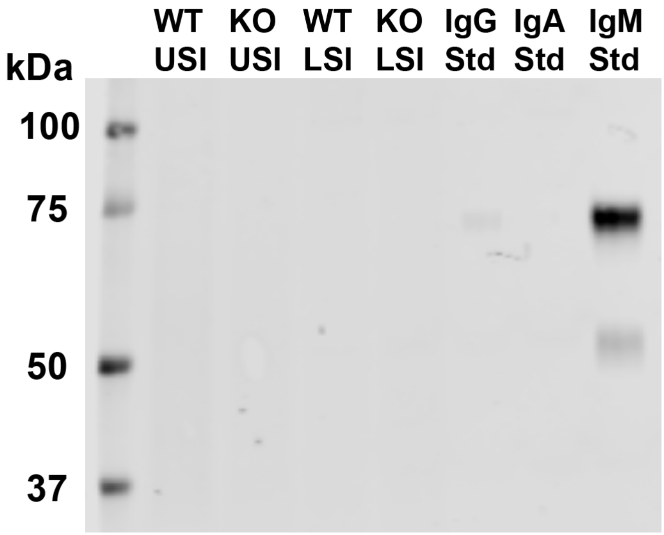


**Figure 6 – figure supplement 2.** Western blots of 30 μm-filtered samples from the small intestine (SI) of wild-type (WT) and Rag1 knockout (Rag1KO) mice. WT USI = WT upper SI; KO USI = KO lower SI; WT LSI = WT lower SI; KO USI = KO upper SI. For detection of IgM, 1:10,000 dilution of Li-Cor IRDye 800CW Goat Anti-Mouse IgM was used. We do not detect IgM in any of the SI samples.

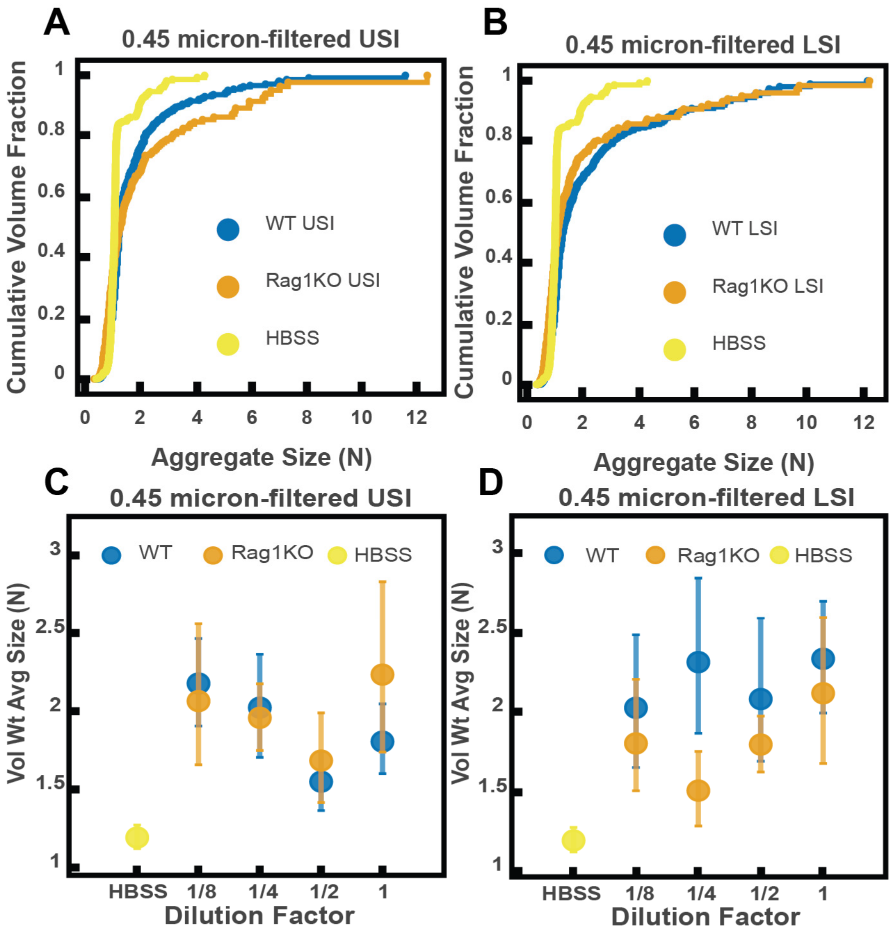


**Figure 6 – figure supplement 3.** *Ex vivo* aggregation in 0.45-µm-filtered luminal fluid from the small intestines (SI) of wild-type (WT) and Rag1 knockout (Rag1KO) mice. (A and B) Volume-weighted empirical cumulative distribution functions (ECDFs) comparing aggregation of the particles in undiluted, 0.45-µm-filtered samples from the upper (A) and lower (B ) SI of two separate groups of WT and immunoglobulin-deficient (Rag1KO) mice to the control (particles suspended in HBSS). Plotted on the vertical axis is the cumulative volume fraction of the total number of particles in solution in an aggregate of a given size. Plotted on the horizontal axis are aggregate sizes in number of particles per aggregate (N). (C and D). Volume-weighted average aggregate sizes (Vol Wt Avg Size) for serial dilutions of 0.45-µm-filtered samples from the upper (C) and lower (D) SI of two separate groups of WT and Rag1KO mice. Volume-weighted average sizes are plotted on the vertical axis in terms of number of particles per aggregate (N). The dilution factor is plotted on the horizontal axis, where a dilution factor of 1 is undiluted and ½ is a two-fold dilution. The control (particles suspended in HBSS) is plotted as a dilution factor of 0. The vertical error bars are 95% empirical bootstrap CI using the bootstrapping procedure described in *Materials and Methods*.

**Table 1. Estimates of physical parameters of polymers from gel permeation chromatography for liquid fractions from the upper small intestine of MUC2 knockout (MUC2KO) and wild-type (WT) mice.**

| **Retention volume (mL)** | **11 to 16** | | **16 to 20** | | **>20** | |
|---|---|---|---|---|---|---|
| **Mouse type** | WT | MUC2KO | WT | MUC2KO | WT | MUC2KO |
| **$M_w$ (kDa)** | 3,560±410 | 5,420±620 | 162±20 | 147±17 | 4.05±0.46 | 2.96±0.34 |
| **$M_w/M_n$** | 1.36 | 1.59 | 2.16 | 2.43 | 3.59 | 10.9 |
| **$R_h$ (nm)** | 49.1 | 45.5 | 6.31 | 5.95 | 1.18 | 1.02 |
| **Fract. Conc. (mg/mL)** | 2.52±0.29 | 1.18±0.13 | 24.6±2.8 | 21.9±2.5 | 88.7±10.1 | 86.0±9.8 |

We calculated values with both dn/dc = 0.185 (for proteins) and dn/dc = 0.147 (pullulan). When the value varied with dn/dc, it is reported in the table as the mid-range values ± the absolute deviation between the two calculated values. $M_w$ = the weight-average molecular weight; $M_w/M_n$ = the dispersity; $R_h$ = hydrodynamic radius; Fract. Conc. = Concentration of a given molecular weight fraction.

**Table 2. Estimates of physical parameters of polymers from gel permeation chromatography for liquid fractions from the lower small intestine of MUC2 knockout (MUC2KO) and wild-type (WT) mice**

| **Retention volume (mL)** | **11 to 16** | | **16 to 20** | | **>20** | |
|---|---|---|---|---|---|---|
| **Mouse type** | WT | MUC2KO | WT | MUC2KO | WT | MUC2KO |
| **$M_w$ (kDa)** | 4,730±540 | 5,180±590 | 219±25 | 155±18 | 13.7±1.6 | 5.93±0.68 |
| **$M_w/M_n$** | 1.24 | 1.80 | 1.91 | 1.84 | 1.88 | 2.03 |
| **$R_h$ (nm)** | 57.0 | 49.2 | 8.45 | 7.58 | 1.89 | 1.35 |
| **Fract. Conc. (mg/mL)** | 3.42±0.39 | 2.36±0.27 | 23.0±2.6 | 22.8±2.6 | 54.8±6.3 | 63.3±7.2 |

We calculated values with both dn/dc = 0.185 (for proteins) and dn/dc = 0.147 (pullulan). When the value varied with dn/dc, it is reported in the table as the mid-range values +/- the absolute deviation between the two calculated values. $M_w$ = the weight-average molecular weight; $M_w/M_n$ = the dispersity; $R_h$ = hydrodynamic radius; Fract. Conc. = Concentration of a given molecular weight fraction.

**Table 3. Estimates of physical parameters of polymers from gel permeation chromatography for liquid fractions from the upper small intestine of immunoglobulin-deficient (Rag1KO) and wild-type WT mice.**

| **Retention volume (mL)** | **11 to 16** | | **16 to 20** | | **>20** | |
|---|---|---|---|---|---|---|
| **Mouse type** | WT | Rag1KO | WT | Rag1KO | WT | Rag1KO |
| **$M_w$ (kDa)** | 1,480±170 | 2,140±250 | 108±12 | 74.2±8.5 | 2.84±0.32 | 1.91±0.22 |
| **$M_w/M_n$** | 1.09 | 1.14 | 2.62 | 2.42 | 1.59 | 1.54 |
| **$R_h$ (nm)** | 31.8 | 39.8 | 4.77 | 2.51 | 1.078 | 0.936 |
| **Fract. Conc. (mg/mL)** | 1.07±0.12 | 1.13±0.13 | 14.3±1.6 | 13.9±1.6 | 66.1±7.6 | 70.5±8.1 |

We calculated values with both dn/dc = 0.185 (for proteins) and dn/dc = 0.147 (pullulan). When the value varied with dn/dc, it is reported in the table as the mid-range value +/- the absolute deviation between the two calculated values. $M_w$ = the weight-average molecular weight; $M_w/M_n$ = the dispersity; Rh = hydrodynamic radius; Fract. Conc. = Concentration of a given molecular weight fraction.

**Table 4. Estimates of physical parameters of polymers from gel permeation chromatography for liquid fractions from the lower small intestine of immunoglobulin-deficient (Rag1KO) and wild-type WT mice.**

| **Retention volume (mL)** | **11 to 16** | | **16 to 20** | | **>20** | |
|---|---|---|---|---|---|---|
| **Mouse type** | WT | Rag1KO | WT | Rag1KO | WT | Rag1KO |
| **$M_w$ (kDa)** | 1,080±120 | 2,490±290 | 66.9±7.7 | 91.6±10.5 | 3.64±0.42 | 3.72±0.43 |
| **$M_w/M_n$** | 1.18 | 1.05 | 1.71 | 1.98 | 2.09 | 1.98 |
| **$R_h$ (nm)** | 34.6 | 47.1 | 4.67 | 4.85 | 1.116 | 1.09 |
| **Fract. Conc. (mg/mL)** | 1.52±0.17 | 1.89±0.22 | 15.8±1.8 | 14.1±1.6 | 49.5±5.7 | 55.1±6.3 |

We calculated values with both dn/dc = 0.185 (for proteins) and dn/dc = 0.147 (pullulan). When the value varied with dn/dc, it is reported in the table as the mid-range values +/- the absolute deviation between the two calculated values. $M_w$ = the weight-average molecular weight; $M_w/M_n$ = the dispersity; $R_h$ = hydrodynamic radius; Fract. Conc. = Concentration of a given molecular weight fraction.

**Table 5: Gel permeation chromatography of Fibersol-2 and pectin in phosphate-buffered saline**

| **Sample** | Fibersol-2 | Pectin |
|---|---|---|
| **$M_w$ (kDa)** | 3.48 | 232 |
| **$M_w/M_n$** | 10.5 | 1.97 |
| **$R_h$ (nm)** | 1.24 | 25.4 |

Both fiber types were analyzed with dn/dc = 0.147 for polysaccharides**.** $M_w$ = weight-average molecular weight; $M_w/M_n$ = the dispersity; $R_h$ = hydrodynamic radius

**Table 6: Estimates of physical parameters of polymers from gel permeation chromatography for liquid fractions from upper small intestine of pectin and Fibersol-2 fed mice**

| **Retention volume (mL)** | **11 to 16** | | **16 to 20** | | **>20** | |
|---|---|---|---|---|---|---|
| **Mouse type** | Pectin | Fibersol-2 | Pectin | Fibersol-2 | Pectin | Fibersol-2 |
| **$M_w$ (kDa)** | 267±31 | 686±79 | 40.0±4.5 | 35.3±4.0 | 1.39±0.16 | 1.67±0.19 |
| **$M_w/M_n$** | 1.50 | 1.08 | 2.15 | 2.64 | 2.45 | 1.48 |
| **$R_h$ (nm)** | 31.8 | N/C** | 5.52 | 2.88 | 0.819 | N/C** |
| **Fract. Conc. (mg/mL)** | 1.62±0.19 | 0.516±0.059 | 9.00±1.03 | 23.3±2.7 | 53.7±6.1 | 77.0±8.8 |

We calculated values with both dn/dc = 0.185 (for proteins) and dn/dc = 0.147 (pullulan). When the value varied with dn/dc, it is reported in the table as the mid-range values +/- the absolute deviation between the two calculated values. $M_w$ = the weight-average molecular weight; $M_w/M_n$ = the dispersity; $R_h$ = hydrodynamic radius; Fract. Conc. = Concentration of a given molecular weight fraction. N/C** denotes values for which the concentration was too low to calculate.

**Table 7: Estimates of physical parameters of polymers from gel permeation chromatography for liquid fractions from lower small intestine of pectin and Fibersol-2-fed mice**

| **Retention volume (mL)** | **11 to 16** | | **16 to 20** | | **>20** | |
|---|---|---|---|---|---|---|
| **Mouse type** | Pectin | Fibersol-2 | Pectin | Fibersol-2 | Pectin | Fibersol-2 |
| **$M_w$ (kDa)** | 282±32 | 1680±190 | 30.2±3.5 | 18.8±2.2 | 1.12±0.13 | 2.32±0.27 |
| **$M_w/M_n$** | 7.37 | 1.64 | 1.70 | 2.78 | 2.89 | 1.14 |
| **$R_h$ (nm)** | 29.0 | 26.4 | 5.28 | 2.16 | 0.724 | 1.06 |
| **Fract. Conc. (mg/mL)** | 2.48±0.28 | 0.839±0.096 | 9.43±1.1 | 53.6±6.1 | 42.7±4.9 | 88.3±10.1 |

We calculated values with both dn/dc = 0.185 (for proteins) and dn/dc = 0.147 (pullulan). When the value varied with dn/dc, it is reported in the table as the mid-range values +/- the absolute deviation between the two calculated values. $M_w$ = the weight-average molecular weight; $M_w/M_n$ = the dispersity; $R_h$ = hydrodynamic radius; Fract. Conc. = concentration of a given molecular weight fraction.

**Table 8. Zeta potential and NMR measurements of PEG-coated particles.** For the zeta potential measurements, each particle solution was 0.1 mg/ml of particles in 1 mM KCl. Measurements were done on a Brookhaven NanoBrook ZetaPALS Potential Analyzer. Three trials were done where each trial was 10 runs each and each run was 10 cycles. Values reported are the average zeta potential for the 30 runs. NMR measurements were performed as described in *Materials and Methods*. Values are estimates of the nanomoles of polyethylene glycol (PEG) per milligrams of particles. To calculate this, we have to assume all the PEG on the surface is a single MW. It is therefore assumed all the PEG on the surface is PEG 5 kDa.

| Surface Modification of PS particles | Zeta potential (mV) | Nanomoles PEG/mg particles |
|---|---|---|
| mPEG 5 kDa | -18.87 ±1.78 | 5.5 |
| mPEG 5 kDa w/ mPEG 1 kDa backfill | -7.66 ±2.12 | 4.6 |
| mPEG 5 kDa w/ mPEG 350 Da backfill | -9.99 ± 1.65 | 4.3 |
| mPEG 5 kDa w/ mPEG 5 kDa backfill | -14.56 ± 1.78 | 4.0 |
| mPEG 2 kDa | -39.59 ± 2.41 | 9.4 |
| Carboxylate-coated (no PEG) | -61.36 ± 12.40 | 0.0 |
